# IN SITU IMAGING OF RETINAL CALCIUM DYNAMICS IN AWAKE ANIMALS

**DOI:** 10.1101/2023.02.11.528145

**Authors:** Yixiang Wang, Ashley Su, Jialin Jin, Daniel Barson, Michael Crair

## Abstract

Mammalian vision starts in the retina. The study of retinal circuits *in vivo* is essential for comprehending retinal neural dynamics under physiological conditions. While several transpupillary retina imaging techniques have been utilized in anesthetized animals, the *in situ* imaging of retinal activity in awake animals has been more difficult to accomplish. These limits have frustrated crucial scientific inquiries, such as how visual processing in the retina is modulated by behavior. In this study, we present novel experimental approaches that stabilize the eye to access *in situ* retinal dynamics with optical techniques in awake mice. Our findings demonstrate that this method can be utilized to: 1) image neural activity in distinct cell types across multiple ages, 2) record meso-scale (e.g. spontaneous retinal waves) or cellular retinal dynamics, 3) study retina functional connectivity *in vivo*, and 4) pharmacologically manipulate retinal activity. We applied these novel approaches to demonstrate that retinal activity is strongly modulated by movement through H1R-dependent histaminergic transmission in vivo, even at the amacrine cell level. These methods are suitable to simultaneously record retinal and brain activity dynamics or to investigate retinal responses to patterned visual stimuli, making accessible fundamental questions about visual processing that have previously been very challenging to achieve.

## Introduction

Electrophysiologic recordings of disassociated optic nerves were reported in the 1920s (Adrian and Matthews, 1928) and 1930s (Hartline, 1938). Since the early 1980s, the isolated retina has become a well-established model to study electrophysiological properties of retinal neurons *in vitro* (Ames and Nesbett, 1981). By tapping into retinal physiology both at single-neuron and circuit levels, *in vitro* studies have greatly extended our understanding of retinal function and computation (Meister and Berry, 1999, Demb and Singer, 2015, Wong et al., 1995, Feller et al., 1996).

Electrophysiologic recordings of retinal ganglion cells (RGC) *in vivo* have also long been established (Kuffler, 1953, Talbot and Kuffler, 1952). Recent advances in calcium indicators and novel microscopy techniques have enabled researchers to image RGC terminals in the thalamus (Liang et al., 2020) and superior colliculus (Ackman et al., 2012, Schroder et al., 2020), demonstrating features of RGC dynamics ranging from synaptic to meso-scale levels. However, imaging retinal outputs does not grant access to neural activity in other retinal cell types, including >60 types of amacrine cells (Yan et al., 2020), ∼15 types of bipolar cells (Shekhar et al., 2016), horizontal cells, photoreceptors, Muller glia cells, or even somata of RGCs (Rheaume et al., 2018). This has frustrated attempts to examine features of intra-retinal functional dynamics *in vivo*.

In the past decade, multiple groups demonstrated the exquisite capability of imaging retinal structures *in vivo* using two-photon (2P) microscopy with adaptive optics (Geng et al., 2012, Qin et al., 2020, Biss et al., 2007, Wahl et al., 2016, Yin et al., 2013, Zhang et al., 2022), including reports of functional imaging of light response (Qin et al., 2020, Yin et al., 2013, Zhang et al., 2022). However, setting up and operating an imaging system with adaptive optics is non-trivial for most research groups. Another approach (Wang et al., 2021) used a slightly modified generic 2P microscope to access the retina and calcium activity with a ratiometric calcium indicator Twitch2b (Thestrup et al., 2014). All these techniques require continuous anesthesia and reported no observations of spontaneous activity. In fact, spontaneous retinal activity is completely abolished even under a minimal amount of anesthesia (Ackman et al., 2012). Moreover, retinal physiology, such as light response, is profoundly impacted by anesthesia (Tomiyama et al., 2016, Nair et al., 2011, Sorensen et al., 2017), highlighting the necessity of examining retinal activity in awake animals.

Here, we report the first *in situ* imaging of retinal activity in awake animals, by both wide-field single-photon microscopy and two-photon microscopy, requiring minimal adaptation of commonly available optical systems. Our method revealed retinal calcium dynamics (Fig. 1, Fig. 2, Supplementary Fig. 1-4) consistent with previous *in vitro* and *in vivo* studies, such as spontaneous retinal waves (Burbridge et al., 2014a, Ackman et al., 2012, Ge et al., 2021a, Gribizis et al., 2019a, Tiriac et al., 2022). We have applied this technique to image mice at different ages (P4-P30) and with *in vivo* retinal pharmacology. Recent studies indicate that arousal might modulate axon boutons of retinal ganglion cells in the thalamus and superior colliculus (Reggiani et al., 2022, Schroder et al., 2020, Liang et al., 2020). In addition to possible presynaptic mechanisms in terminal brain regions, previous reports showed that histaminergic retinopetal axons branch extensively in the ganglion cell and inner plexiform layers (Greferath et al., 2009; Gastinger et al., 2006). In vitro results indicate that efferent projections from the hypothalamus can modulate retinal ganglion neurons (Warwick et al., 2022). Utilizing our unique setup to image the retina *in situ* in awake animals, we demonstrated that some amacrine cells were substantially modulated by movement mediated through the H1 receptor during early development, suggesting that the retina receives ongoing efferent feedback from the brain starting at an early age.

**Fig. 1.**
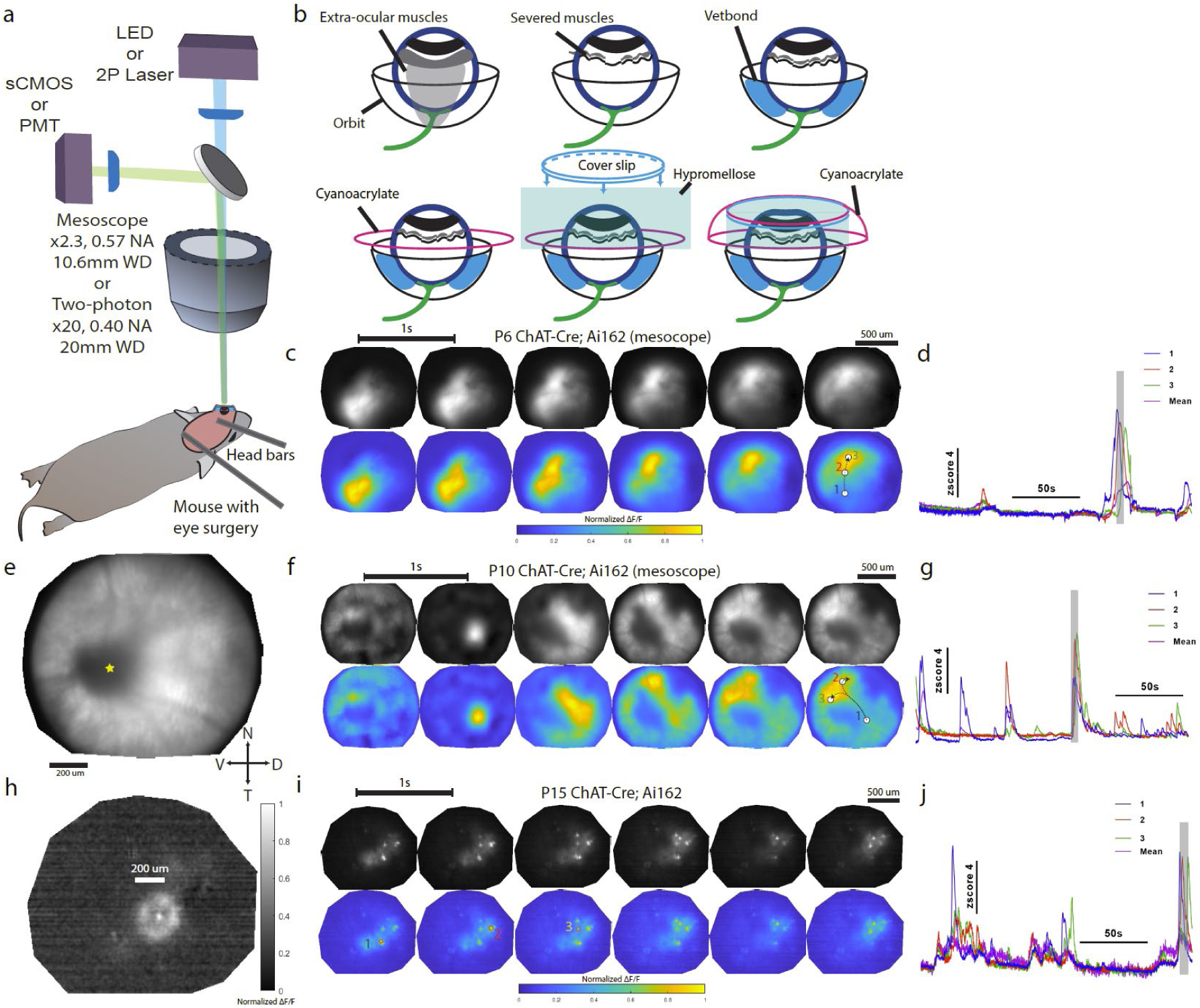
*In situ* imaging of retinal calcium dynamics in awake animals. a. A simplified schematic showing the experiment setup of *in vivo* retina imaging in awake animals with single-photon (1P) wide-field or two-photon (2P) microscopes. b. Major steps of the eye surgery. For details see Method. c. Example montages of mesoscopic retina activity (retinal waves) from a P6 ChAT-Cre; Ai162 animal. Colormaps: Montage interval = 1 second. ΔF/F values were normalized to [0, 1] for plotting. Top row: grayscale (raw fluorescence); Bottom row: parula (MATLAB). White dots denote three pixels/ regions of interest (ROIs) corresponding to the activity traces in panel d. d. Z-scored activity traces from three pixels/ ROIs denoted in panel c. Shaded area corresponds to the period depicted in c. e. An example image of the field of view (FOV) generated by averaging over 3000 frames. Yellow star: optic disc. f. Example montages of mesoscopic retina activity (retinal waves) from a P10 ChAT-Cre; Ai162 animal. Colormaps: Montage interval = 1 second. ΔF/F values were normalized to [0, 1] for plotting. Top row: grayscale (raw fluorescence); Bottom row: parula (MATLAB). White dots denote three ROIs corresponding to the activity traces in panel g. g. Z-scored activity traces from three pixels/ ROIs denoted in panel f. Shaded area corresponds to the period depicted in f. h. An example of an active starburst amacrine cell from the same animal as in i. Colorbar: gracale representing ΔF/F values normalized to [0, 1] i. Example montages of sparse retina activity from a P15 ChAT-Cre; Ai162 animal. Colormaps: Montage interval = 1 second. ΔF/F values were normalized to [0, 1] for plotting. Top row: grayscale (raw fluorescence); Bottom row: parula (MATLAB). Red dashed circles denote three ROIs corresponding to the activity traces in panel j. j. Z-scored activity traces from three pixels/ ROIs denoted in panel i. Shaded area corresponds to period depicted in in i.

**Fig. 2.**
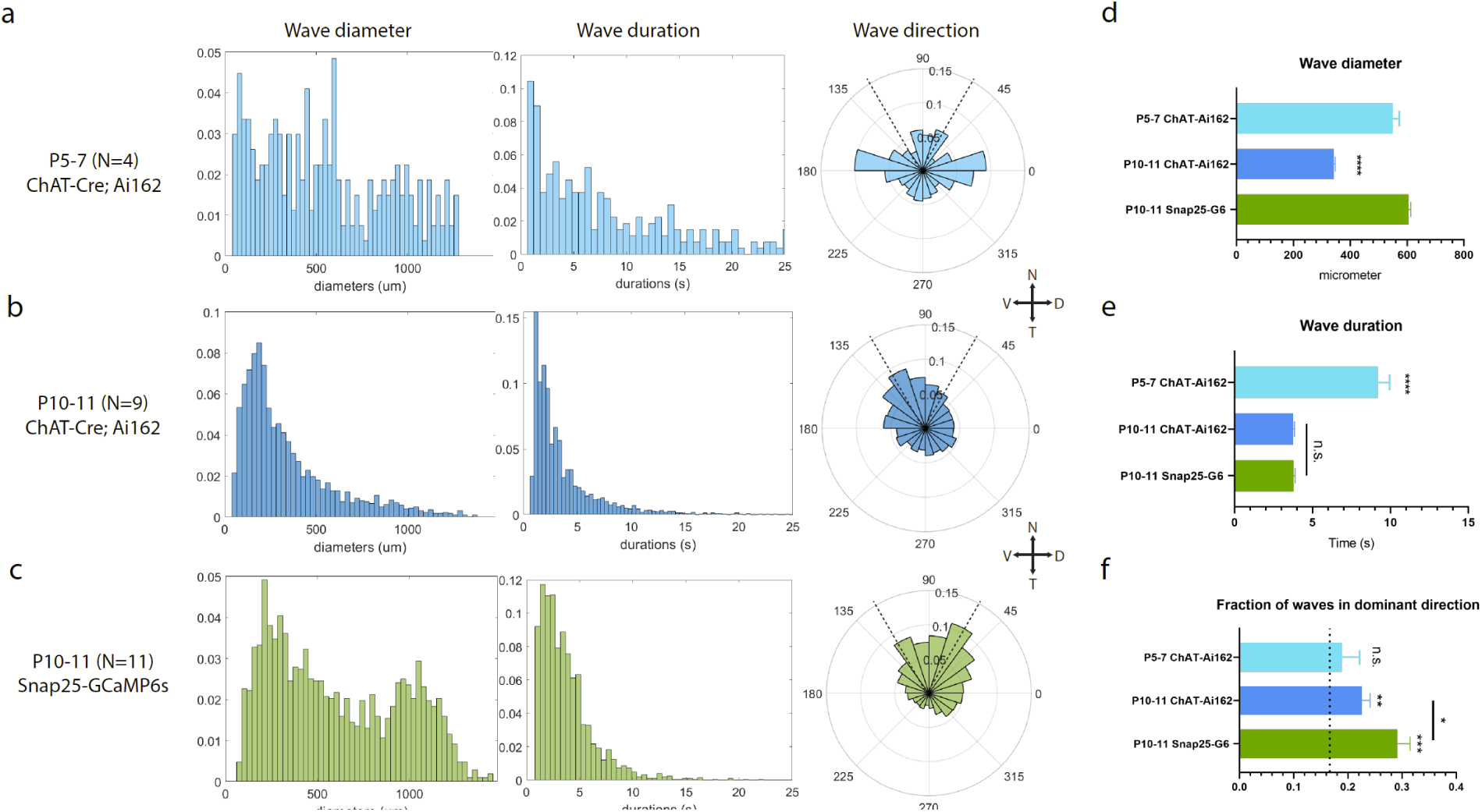
Spatiotemporal properties and directionality of in situ retinal waves. a. Spatiotemporal properties of stage II waves in P5-P7 ChAT-Cre; Ai162 animals (N = 4). Left: wave diameter distribution (normalized histogram, x-axis unit: um); Middle: wave duration distribution (normalized histogram, x-axis unit: seconds); Right: frequency distributions of waves directions (normalized polar histogram), 0 degree corresponds to the dorsal direction. Crossed double-headed arrows indicate cardinal directions (N: nasal; V: ventral; D: dorsal; T: temporal). Dashed black lines indicate the 60° around the dominant direction that was used for further analyses. b. Spatiotemporal properties of stage III waves in P10-P11 ChAT-Cre; Ai162 animals (N = 9). Panel layout similar to a. c. Spatiotemporal properties of stage III waves in P10-P11 SNAP25-GCaMP6s animals (N = 11). Panel layout similar to a. d. Retinal waves from ChAT-Cre; Ai162 animals at P10-P11 (stage III) were significantly smaller than those from SNAP25-GCaMP6s animals at the same age and/or those from ChAT-Cre; Ai162 animals at P5-P7 (stage II). e. Retinal waves from ChAT-Cre; Ai162 animals at P5-P7 (stage II) had significantly longer durations than those from SNAP25-GCaMP6s animals at P10-P11 (stage III) and/or those from ChAT-Cre; Ai162 animals at P10-P11. f. Proportion of wave propagation directions within 60° surrounding the dominant direction (temporal-nasal). Proportions of wave propagation directions within 60° surrounding the dominant direction were significantly larger than random (indicated by the dashed line, one sample t-test) for P10-P11 SNAP25 and ChAT waves. Besides, a significantly higher fraction of P10-P11 SNAP25 waves were in the 60° range in compared to P10-P11 ChAT waves. n.s. p > 0.05, ∗ p < 0.05, ∗∗ p < 0.01, ∗∗∗ p < 0.001, **** p < 0.0001, two-tailed unpaired t-test with Welch’s correction.

## Results

### In situ Imaging of Retinal Calcium Dynamics in Awake Animals

The challenges associated with imaging retinal activity in awake animals are twofold. Firstly, the constant movement of the eye presents difficulties in obtaining clear and stable images of the retina as even slight eye movements can result in significant image blur. Secondly, the occurrence of cataracts, caused by factors such as dehydration or other stressors that result in lens opacity, further complicates imaging efforts.

To address these challenges, a new procedure has been developed that significantly reduces eye movement through surgical intervention and enables extended imaging sessions with minimal lens cloudiness. In brief: mice were anaesthetized with isoflurane throughout the surgery; Anaesthetized mice were also given analgesics and then placed on a ball stage that can tilt from 0-90 degree; A piece of scalp (including the face skin near the left eye) was removed, and head bars were positioned to the dorsal surface and behind the occipital skull; Cyanoacrylate was used to glue the head bars to the head; Extra-ocular muscles were carefully severed without damaging major periorbital blood vessels; a small amount of tissue-adhesive Vetbond (3M) was injected behind the eyeball to stick the back of the eye to surrounding orbit/ eye socket; Cyanoacrylate was applied to surface of exposed skull near the eyeball to further stiffen and stabilize the surrounding region; Atropine was given topically to dilate the pupil; Hydrophilic Hypromellose of high viscosity was dropped to completely submerge the cornea and keep it hydrated throughout imaging sessions (critical to keep ocular media clear); A coverslip was placed onto the Hypromellose droplet and was pushed down against the cornea to create a relatively flat surface; The structure was then sealed with additional cyanoacrylate to enclose the submerged eyeball underneath the cover slip (Schematics: Fig. 1b; Photos: Supplementary Fig. 1a; see Methods and Supplementary Movie 1 for details).

After the animals recovered from anesthesia, the ball stage was tilted and placed under a single-photon (1p) mesoscopic microscope or a two-photon (2p) microscope for imaging (Fig. 1a). Under the one-photon mesoscopic microscope, the optic disc and surrounding vasculature could be seen when the ocular media was clear (Fig. 1e). We successfully recorded mesoscopic retinal dynamics (spontaneous retinal waves) in animals prior to eye-opening (Fig. 1, Supplementary Fig. 1-2). Specifically, we observed stage II retinal waves at P6-P7 (see example montages and fluorescent traces in Fig. 1 c-d) and stage III retinal waves at P10-P11 (Fig. 1f-g) selectively in cholinergic neurons (presumably starburst amacrine cells) by imaging ChAT-Cre; Ai162 animals (Daigle et al., 2018). We also observed robust stage III retinal waves at P10-P11 pan-neuronally using SNAP25-GCaMP6s animals (Fig. 2 and Supplementary Fig. 4), as well as selectively in presumptive bipolar cels and photoreceptors (Johnson et al., 2007) using Vglut1-Cre; Ai162 animals (Supplementary Fig. 1b). After eye-opening, spontaneous retinal waves ceased, but sparse firings still occurred (Fig. 1i-j), which allowed us to access retinal dynamics at the cellular level. We show an example of a single starburst amacrine cell fluorescing with its soma at the center and a halo (presumably its dendritic arbor) around the cell body (Fig. 1h). We observed no spontaneous calcium dynamics in mice under anesthesia (0.5% isoflurane, P8-P10, N = 3, data not shown).

In the single-photon imaging setup, we used 4-8mW (depending on the opacity of ocular media) blue LED peaked at ∼460nm (incident intensity at cornea surface was approximately 1-2mW/mm^2^) to excited GCaMP. We were concerned that this excitation light might saturate rods and obscure ongoing retinal activity despite adaptation of the photoreceptors (Tikidji-Hamburyan et al., 2017). However, we demonstrated that the retina could still respond to violet light (∼3mW violet LED peaked at ∼385nm) on top of constant blue illumination light (∼8mW blue LED), suggesting the retina was not saturated under this condition (Supplementary Fig. 1c).

As mouse opsins are less sensitive to far-red/infra-red lights (Wang et al., 2011), we also explored far-red indicators with red-shifted illumination (∼635nm) or GCaMP6s with two-photon illumination (920nm). Specifically, we tested four far-red indicators: FR- GECO1c, HaloCaMP1b (with *in vivo* injection of JF635, see method for details), NIR- GECO2G, and iGECI (Dalangin et al., 2020, Deo et al., 2020, Qian et al., 2020, Shemetov et al., 2021) for single-photon mesoscopic imaging (supplementary Fig. 2c). Far-red indicators were generally dimmer than GCaMP despite better tissue penetration by red-shifted emissions (Shcherbakova, 2021). In our experiments, with the maximum LED power (∼90mW, ∼635nm), only FR-GECO1c exhibited detectable fluorescent signals despite a worse signal-to-noise ratio and faster bleaching relative to GCaMP6s (supplementary Fig. 2a-b).

Two-photon imaging allows the acquisition of GCaMP signals at an excitation wavelength of >920nm. We therefore aimed to use a conventional two-photon microscope with no adaptations to image GCaMP in the retina in awake animals (see Methods for details). In brief, we conducted intravitreal injection of AAV2.1-hsyn-jGCaMP7b (Dana et al., 2019) at P0 in ChAT-Cre; Ai9 animals (Madisen et al., 2010), and imaged at later ages (>P10). Because of the optical sectioning effect of 2P laser, precisely locating the focal plane to the retina is critical. Expression of tdTomato could greatly facilitate this process due to its high brightness (Soleja et al., 2018), although jGCaMP7b (providing the highest baseline brightness among all existing GCaMPs) alone was sufficient for locating the retina with a longer search time. We observed spontaneous calcium activity in a P13 animal expressing jGCaMP7b (Supplementary Fig. 2 d-e) using 920nm laser at moderate power (50mW). To acquire better imaging quality, we explored raising the laser power to up to 100mW and could still observe activity after 30min of imaging (data not shown). However, higher power levels likely would induce permanent damage to the retina (Wang et al., 2021), due to phototoxicity and heating (Khan et al., 2015, Nourhashemi et al., 2016, Podgorski and Ranganathan, 2016).

To gain a more quantitative perspective on how each approach might activate photoreceptors (1P imaging of GCaMP with low-power blue LED, 1P imaging of FR- GECO1c with high-power far-red LED, and 2P imaging of GCaMP with 920nm laser), we built a mathematical model considering the following factors (Supplementary Fig. 2g, see Methods for details): 1) incident LED/laser power at the objective (data from manufacturers, translated to photon influx), 2) bandpass filtering (data from the manufacturer), 3) transmittance of ocular media (Lei and Yao, 2006b), and 4) spectral sensitivity of mouse opsins (Lamb, 1995a). Compared to the activation level induced by a low-irradiance blue LED (BDX, X-Cite with power at 5%, see Methods for details), relative activation of mouse rhodopsin by high-irradiance far-red light was ∼15.9%, whereas instant activation by 2P laser (920nm) was ∼1.36% (during the ∼150ns laser dwelling time, or ∼5.2×10^-8 on average). In summary, imaging the retina with a far-red LED or infra-red laser could potentially reduce activations of photoreceptors, but at the cost of image quality.

Despite these caveats, using our 2P imaging data, we uncovered distinct functional connectivity domains in the retina (Supplementary Fig. 3). We conducted seed-based correlation analysis and placed references seeds at three different locations that were >250um apart (supplementary Fig. 3a). The three corresponding correlation maps were then binarized at a threshold = 0.4. Less than 3% of pixels of domain 1 and 2 colocalized, in contrast to 80% overlap of pixels between domains 2 and 3 (supplementary Fig. 3b), suggesting the presence of distinct functional connectivity modules in the retina in awake animals.

### Spatiotemporal properties and directionality of in situ retinal waves

Despite first being characterized *in vitro* (Meister et al., 1991), retinal waves are only observed *in vivo* in unanesthetized animals (Ackman et al., 2012). Previous *in vivo* studies investigated retinal waves by imaging RGC axons in the superior colliculus (Ge et al., 2021b, Gribizis et al., 2019b, Xu et al., 2016, Burbridge et al., 2014b, Ackman et al., 2012). Using the *in situ* retina imaging technique, we recorded mesoscopic spontaneous retinal waves in different cell types within the retina and under various conditions (Fig. 1c- j, Supplementary Fig. 1b, Figure 2, Supplementary Fig. 4, and Supplementary Movie 2).

From previous recordings of retinal ganglion cell (RGC) axons in the superior colliculus (SC), we knew that retinal waves propagate in the temporal-to-nasal direction at P8-P11 (Ge et al., 2021a). We recapitulated this finding using SNAP25-GCaMP6s mice (Fig. 2c), which express calcium indicators in all retinal neurons including RGCs. However, given the critical role of starburst amacrine cells (SAC) in mediating wave directions, we wondered if wave properties could differ if calcium signals were confined to SACs. By using ChAT-Cre; Ai162 mice, we found that retinal waves in ChAT+ neurons, referred to as “ChAT waves” thereafter, were significantly larger and lasted longer at stage II (P5-7) compared to stage III (P10-P11), as demonstrated in Fig. 2d-e. These results align with previous studies that reported larger and slower stage II waves in comparison to stage III waves (Gribizis et al., 2019a, Arroyo and Feller, 2016). However, retinal waves in SNAP25-GCaMP6s animals (“SNAP25 waves” thereafter) at stage III had similar durations to stage III ChAT waves (Fig. 2e). Besides, stage III ChAT waves also showed directional bias but with a slight shift toward the ventral direction (Fig. 3b). As a result, stage III SNAP25 waves were more concentrated in the dominant direction (higher fraction of waves within the 60° range around the nasal cardinal axis) than ChAT waves (Fig. 2f).

**Fig. 3.**
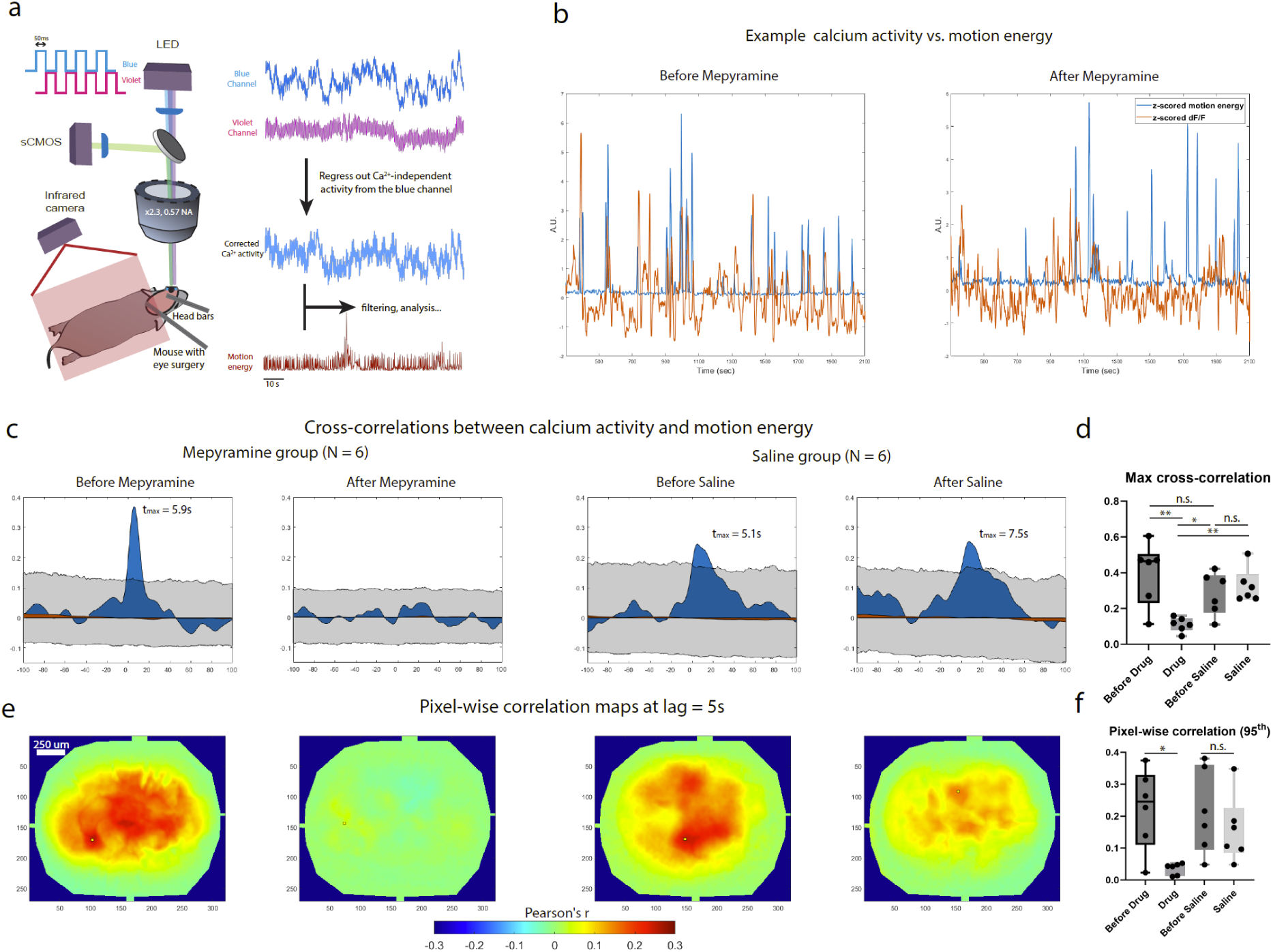
Movement modulates retinal activity through an H1R-dependent mechanism. a. Experimental setup of *in vivo* retina imaging in awake animals with a single-photon microscope and movement recording with an infrared camera. Movies were acquired with interleaved frames of violet (395 nm) and blue (470 nm) illumination at 10Hz. Example traces on the right (top to bottom) shows 1) mean z-scored ΔF/F over all pixels in the field of view from the blue channel (470nm illumination), 2) mean z-scored ΔF/F over all pixels in the field of view from the violet channel (395nm illumination), 3) mean corrected z-scored ΔF/F overall all pixels in the field of view after regressing out the violet channel activity (Ca2+-independent) from blue channel activity, and 4) z-scored motion energy of mouse body movement. b. Example traces of calcium activity against simultaneously recorded motion energy before and after injection of 200nL 500uM mepyramine in the same animal. Orange trace: calcium signal was calculated by averaging ΔF/F values across all pixels in the field of view. Blue trace: motion energy was computed by accumulating pixel-wise illuminance changes between neighboring frames of body movements captured by the infrared camera. Mean calcium activity and motion energy traces were z-scored to display at the same scale. See Method for details. c. Average cross-correlation between mean calcium activity and motion energy (blue) across 6 animals (P12-P16) in the mepyramine group (1^st^ and 2^nd^ panels from left to right: before and after drug injection in the same animals, and across 6 animals (P12-P16) in the saline group (3^rd^ and 4^th^ panels from left to right: before and after saline injection in the same animal). Grey shade shows the 2.5th to 97.5th percentile of circularly permutated (time-shifted) data. Maximum cross-correlations were found at t_max_ > 5s before mepyramine injection and before/after saline injection (positive t_max_ indicates the amount of time that calcium activity shifted to trail movement). The orange area denotes the result after random temporal permutation. See Method for details. d. Quantification of max cross-correlations for each animal. Mepyramine injection significantly reduced maximum cross-correlations compared to controls. e. Average correlation maps across multiple animals illustrating pixel-wise correlation between calcium activity (ΔF/F) and motion energy at time lag = 5s (calcium activity trailing movement). The order is the same as in panel c. Small yellow squared denote the pixels that have the 95^th^ percentile largest correlation. f. Quantification of the 95^th^ percentile (largest) correlation for each animal. Mepyramine injection significantly reduced the correlations compared to controls. n.s. p > 0.05, ∗ p < 0.05, ∗∗ p < 0.01, ∗∗∗ p < 0.001, **** p < 0.0001, two-tailed unpaired t-test with Welch’s correction. For all Box plots in this study: hinges: 25 percentile (top), 75 percentile (bottom). Box whiskers (bars): Max value (top), Min value (bottom). The line in the middle of the box is plotted at the median.

To disrupt spatiotemporal properties of retinal waves *in vivo*, we re-anesthetized previously imaged animals and injected an inhibitory antagonist cocktail intravitreally (200uM Gabazine + 100uM Strychnine + 2mM TPMPA, to block GABA-A, glycine, and GABA-C receptors). Waves after the cocktail application had significantly larger diameter and longer duration (Supplementary Fig. 4 e-f), presumably due to reduction of inhibition in the retina, but continued to exhibit a propagation bias in the temporal-to-nasal direction.

### Movement modulates in situ retinal activity through the histamine H1 receptor

Previous studies (cite Schroeder and Liang) have demonstrated modulation of RGC axon activity by movement but the effects of movement on the activity of RGCs and other cell types within the retina remains unknown. As our method enables cell type-specific measurement of retina activity *in situ* in awake animals, we investigated whether ChAT-positive neurons in the retina are modulated by movement by single-photon mesoscopic imaging of ChAT-Cre; Ai162 mice at P12-P16 with concurrent measurement of movement using an infrared camera. We chose this age window as propagating retinal waves cease but bursts of spontaneous activity can still be observed (Supplementary Movie 3).

We measured correlations between retina calcium activity and movement (quantified as motion energy of body movements, see Methods for details). Although the mouse pupil was dilated throughout the imaging and surgery procedures stabilized the eyeball, substantial body movements sometimes caused movements of the eye that obscured imaging (such as squeezing along the z-axis). In addition, hemodynamic changes can confound fluorescent measurements of neuronal activity. To reduce these artifacts, we illuminated the retina with interleaved blue and violet light, following a well-established protocol (Allen et al., 2017, Barson et al., 2020). GCaMP emissions under violet illumination are largely isosbestic / Ca^2+^-independent (Lerner et al., 2015). We regressed out this isosbestic signal from the blue-light-illuminated trace to remove concurrent motion and hemodynamic-related artifacts, rendering only Ca^2+^-dependent signal (Fig. 3a). Motion energy was computed based on pixel-wise illuminance differences between neighboring frames of body movement (see Methods for details). We then computed the cross-correlation between the mean calcium activity (averaging across all pixels in the field of view, after regressing out the isosbestic component) and the concurrent movement. Example ΔF/F and motion energy traces are shown in Fig. 3b. We discovered significant positive cross-correlations peaked at ∼5s lags (calcium activity trailing movement, Fig. 3c and Supplementary Fig. 5e) compared to permutated data. To understand spatial patterns of such modulation at the single-pixel level, we plotted correlation maps with calcium activity trailing movement at a lag=5s (Fig. 3e).

A previous study reported top-down histaminergic modulation of retina activity (Warwick et al., 2022)) mediated by the H1 receptor (H1R). To test whether movement modulates retina activity through H1R, we injected H1R blocker mepyramine (200nL, 500uM) to the eye and performed repeat in situ retinal calcium imaging. Mepyramine significantly reduced the cross-correlation between retina calcium activity and movement (Fig. 3c-d, P12-P16, N = 6). Similarly, when examining pixel-wise correlation maps at a lag = 5s (Fig. 3e), mepyramine abolished the overall correlation (Fig. 3e-f). In the saline injection group (P12-P16, N = 6), however, no such effect was observed. (Fig. 3c-f, N = 6). Mepyramine did not affect the frequency and peak amplitude of calcium activity (measured by ΔF/F peaks, Supplementary Fig. 5), and differences in calcium activity levels did not affect correlational outcomes (Supplementary Fig. 6). Therefore, our results indicate that movement modulates retina activity through the histaminergic H1 receptor.

## Discussion

We describe *in situ* wide-field and two-photon imaging of retinal activity in awake animals for the first time. Advantages of our approach include: 1) implementation using commonly available, turn-key wide-field epifluorescence or two-photon microscopes; specifically, it does not require adaptive optics or customized contact lenses for rodents. 2) feasibility in mice across various ages and sizes (P4 to adult mice), 3) compatibility with pharmacology experiments, and 4) compatibility with simultaneous “dual-imaging” of retinal and brain activity (Barson et al., 2020).

One caveat of this method is that the continuous blue excitation light (∼460nm) used in single-photon imaging sessions may saturate rod photoreceptors. However, the retina was still responsive to pulsed violet light during imaging sessions with contuous blue excitation light at intensities up to 2 mW/mm^2, which was the maximum assessed in our study (Supplementary Fig. 1c). To reduce potential single-photon and two-photon phototoxicity to the retina, we kept the blue LED power at the minimum level necessary for consistent recording of spontaneous activity. Spontaneous retinal dynamics could still be observed after hours of continuous acquisition under all conditions in this study without noticeable disruptions to ongoing activity.

To reduce the impact of blue excitation light on retinal activity, we tried to block signaling from photoreceptors to ON and OFF bipolar cells with a combination of L-AP4 (200uM) and UBP 310 (500uM), informed by previous *in vitro* experiments (Tiriac et al., 2018). Responses to blue excitation light were reduced by 50% when measured 30 minutes after injection (minimum time for animals to recover from anesthetics), but recovered to baseline amplitudes after 1 - 1.5 hours (data not shown). We also sought to mitigate this issue by using far-red calcium indicators or two-photon imaging. Among the four far-red indicators tested, only FR-GECO1c (Supplementary Fig. 2) yielded detectable wide-field fluorescent signals for *in situ* retina imaging, while the rest iGECI and NIR- GECO2G yieleded detectable signals in cortex but not in the retina (data not shown). Based on our tests and a previous review of far-red indicators (Shcherbakova, 2021), we concluded that two factors might be important to predict whether a far-red indicator is suitable for *in situ* retina imaging: 1) maximum brightness relative to EGFP (>25%), and 2) dynamic range (>15X). We also imaged retinal dynamics *in situ* with a conventional two-photon microscope (Supplementary Fig. 2) using a relatively high laser power (50mW-100mW). Retina activity could be recorded for at least one hour at 100mW, but damage to the retina was observed at higher powers (data not shown). To quantify the impact of different imaging schemes, we modeled the relative activations of rhodopsin (Supplementary Fig. 2). The result suggested that imaging FR-GECO1c (single-photon LED) or imaging GCaMP with a two-photon infrared laser were potentially beneficial for reducing activation of photoreceptors. However, these modifications resulted in reduced signal-to-noise ratio and image quality and might cause other issues (FR-GECO1c bleached much faster than GCaMP, while the two-photon infrared laser might cause tissue heating) suggesting thatfurther optimization is necessary to leverage these modifications to reduce photic stress to the retina.

Using our imaging approach, we quantified spatiotemporal properties and directionality of *in situ* retinal waves (Fig. 2 and Supplementary Fig. 4). In general, we observed smaller, briefer, and directionally biased stage III retinal waves compared to stage II waves, consistent with previous reports (Ge et al., 2021a, Gribizis et al., 2019a). These experiments yielded three novel observations. First, stage III ChAT waves were smaller than stage II ChAT waves (Fig. 2d), consistent with previous reports of smaller, briefer, and directionally biased stage III retinal waves relative to stage II waves (Elstrott and Feller, 2010, Maccione et al., 2014, Gribizis et al., 2019a, Ge et al., 2021a). Second, stage III ChAT waves were smaller than stage III SNAP25 waves (Fig. 2d). This suggested that stage III ChAT waves (from cholinergic starburst amacrine cells) might be a component of SNAP25 waves (pan-neuronal expression of GCaMP6s in all retinal neurons). Third, directional bias persisted after pharmacologically blocking major inhibitory receptors using a cocktail of gabazine, strychnine, and TPMPA (Supplementary Fig. 4). In previous work from our group (Ge et al., 2021a), gabazine alone was shown to abolish directional bias. We speculate that further reduction of inhibition in the retina using the expanded cocktail unmasks the default “initiation bias” in the temporal retina, which was also observed in late stage II waves (Ge et al., 2021a). Alternatively, evaluating the developing retina from a dynamical system perspective (Matzakos-Karvouniari et al., 2019) might explain the discrepancy, as the system could be pushed to different regimes given different excitation and inhibition levels.

Another caveat is that the blue illumination used to image GCaMP may alter wave properties, as constant light was shown to impact stage III waves *in vitro* (Tiriac et al., 2018). However, previous studies indicated that photoreceptors were insensitive to blue lights before P8 (Bonezzi et al., 2018), suggesting the impact of external illumination on early stage II waves should be limited.

Our method offers a unique opportunity to study the *in situ* retinal physiology in awake animals, allowing us to correlate retinal dynamics with movement, as recent studies imaging RGC boutons in the superior colliculus (Schroder et al., 2020) and thalamus (Liang et al., 2020) demonstrated that retinal outputs are modulated by movement and arousal. In our current setup, the pupil was dilated by atropine sulfate throughout imaging sessions, and the eye was exposed to a constant illumination pattern (interleaved blue and violet light at 10Hz). Therefore the illuminance level at the retina was stable and not contingent on pupil dynamics. We observed significant cross-correlation (peaking at lag ∼5s) between movement and calcium dynamics in ChAT-positive neurons (Fig. 3) and speculated that retinopetal projections from the brain might play an important role in this slow modulation. In rodents, these projections originate primarily in the hypothalamus and the raphe nucleus, are histaminergic and serotonergic, respectively (Gastinger et al., 2006), and could have slow and long-lasting effects (Celada et al., 2013, Frazao et al., 2011). Recent in vitro evidence also suggests that top-down projection from the hypothalamus modulates retina activity through H1R (Warwick et al., 2022), possibly through effects on.horizontal cells and multiple types of amacrine cells (Frazao et al., 2011, Yu et al., 2011). In accordance with these studies, we demonstrated that H1R antagonist mepyramine could significantly reduce the cross-correlation between movement and retinal calcium dynamics (Fig. 3), indicating histaminergic transmission mediates movement-related top-down modulation. As we selectively expressed calcium indicators in ChAT+ retinal neurons (presumably starburst amacrine cells), these results collectively suggest that multiple cell types in the retina can be potentially modulated by histaminergic efferent projections.

## Methods

### Transgenic Models

SNAP25-GCaMP6s animals: B6.Cg-SNAP25tm1Hze/J (JAX#025111) animals that have pan-neuronal GCaMP6s expression were used widely in this study. ChAT-Cre; Ai162 animals: Homozygous ChAT-Cre animals (JAX #018957) were crossed to homozygous TIT2L-GCaMP6s-tTA2 animals (Ai162; JAX #031562) to generate offspring that express GCaMP6s in ChAT+ neurons. ChAT-Cre; Ai9 animals: Homozygous ChAT-Cre animals (JAX #018957) were crossed to homozygous Rosa26-CAG-LSL-tdTomato animals (Ai9; JAX # 007909) to generate offspring that express tdTomato in ChAT+ neurons. Vglut1-Cre; Ai162 animals: Homozygous Vglut1-Cre animals (JAX # 023527) were crossed to homozygous Ai162 animals (JAX #031562) to generate offspring that express GCaMP6s in Vglut1+ neurons. WT: C57BL/6J (JAX # 000664).

### Animals Usage

Animals of both sexes were used in this study. Animal care and use followed the Yale Institutional Animal Care and Use Committee (IACUC), the US Department of Health and Human Services, and institution guidelines. In Fig. 1 and Fig. 2, ChAT-Cre; Ai162, animals between P5-P15 were used for in *situ* retina imaging for spatiotemporal and correlation analysis at different ages. In Supplementary Fig.1 P10 WT and P13 ChAT-Cre; Ai9 mice were used for retina imaging. In Supplementary Fig. 2, P10 Vglut1-Cre; Ai162 animals were used for retina imaging. In Fig. 2 and Supplementary Fig. 4, SNAP25-GCaMP6s animals were used. In Fig. 3, ChAT-Cre; Ai162 animals between P12-P16 were used.

### Surgery For *In Vivo* Imaging

#### Surgery for *in situ* retina imaging in awake animals

Animals were anesthetized with 2-3% isoflurane in oxygen via a nose cone and placed on a heating pad set to 37°C (HTP-1500 Adroit). Carprofen was administered subcutaneously (5 mg/kg), and local anesthesia was provided topically using lidocaine (0.5%). Approximately 2-mm of the scalp was then removed to expose the skull, which was then dried with a surgical swab sprayed with 70% isopropyl alcohol. Two steel head posts were then attached to the skull with cyanoacrylate glue: one beneath the occipital portion of the skull to provide support and the other resting on the lambda point. These posts were attached to an articulating base ball stage (SL20, Thorlabs) that allowed for rapid angular positioning. The eyelid and approximately 0.5mm of surrounding skin were then carefully removed from the eye. Initial drops of atropine sulfate (1%, made from Millipore Sigma CAS: 5908-99-6) were administered to dilate the pupil, and subsequent drops of Hypromellose (2.5% dissolved in 1XPBS, made from Millipore Sigma CAS: 9004-65-3) were administered frequently to maintain a moist environment. The extra-ocular muscles were cut with microdissection scissors and approximately 10 uL of Vetbond was carefully injected behind the eyeball to fix the eyeball to the orbit. The eyeball was further fixed from above with a layer of cyanoacrylate, covering the exposed outer area of the eyeball and surrounding skull, ensuring that the pupil remained clear and unobstructed. The exposed pupil was then covered with a 2.5% solution of Hypromellose and then covered with a round 5mm glass coverslip. This glass coverslip was glued to the exposed skull with cyanoacrylate. After surgery, the animal was left to recover on the heat pad with an oxygen supply for 90 minutes before imaging.

#### Acute pharmacology and reinstallation of coverslip

After data acquisition before eye injection, animals were re-anesthetized with 1-1.5% isoflurane. Articulating ball stage was adjusted to allow better angular positions to perform eye injections. The previously installed coverslip was removed carefully with forceps, and additional drops of atropine sulfate (1%) and Hypromellose (2.5%) were applied to ensure submergence of the cornea surface. For experiments related to Fig. 2: cocktail solution (200uM Gabazine + 100uM Strychnine + 2mM TPMPA. CAS#: 104104-50-9, 1421-86-9, 182485-36-5, respectively) was loaded to a glass pipette attached to a Nanoject III (Drummond Scientific Co). The eye was punctured by the sharp glass pipette, and 250 or 300nL (250nL for animals less than 5g) of cocktail solution was injected intraocularly. For experiments related to Fig. 3: Mepyramine solution (200nL 500uM. CAS# 59-33-6) was used. 2.5% Hypromellose was supplemented after the injection, and a new 5mm round coverslip was sealed with cyanoacrylate. Animals were allowed at least 30min to recover (with oxygen delivered) before imaging sessions.

### *In situ* retina imaging in awake animals

#### Imaging with a wide-field single-photon microscope

Schematics of wide-field imaging apparatus are shown in Fig. 1a. The articulating ball stage was tilted to ensure that the tangent surface of the cornea was perpendicular to incoming excitation lights. A sCMOS camera (pco.edge, PCO) coupled to a Zeiss AxioZoom V16 microscope with an objective PlanNeoFluar Z 2.3x/0.57 FWD 10.6mm was used for imaging acquisition. During imaging sessions, pups were wrapped with cotton gauze during data acquisition as described in (Gribizis et al., 2019a). Magnifications varied based on the sizes of dilated pupils in different animals. Each recording session contained continuously acquired movies for >40 mins. The retina was allowed 10min under different but consistent illumination conditions (constant blue, far-red LED, alternating blue/ violet, or infrared laser) for adaptation: the first 10min of recorded activity was not included in data analysis/ or used for illustration. For Fig. 1-2 and related supplementary figures, data was acquired with blue LED illumination (BDX, X-Cite XLED1 LED) at 10 Hz. For Fig. 3, interleaving frames were illuminated with blue or violet (BDX, UVX, X-Cite XLED1 LED) lights at 20Hz. LED switching and the infra-red camera were triggered by sCMOS camera frames (acquisition rate: 20Hz, exposure time: 49ms). sCMOS camera and alternating LED illuminations were synchronized by a Master-8 (AMPI) stimulator. All timing data was collected on a Power3 DAQ and Spike2 software (Cambridge Electronic Design). **Filter sets and incident intensities**: For blue-illuminated (BDX) or blue and violet (BDX and UVX) interleaving illumination, a T495lpxr dichroic mirror was used with an ET525/50m emission filter (Chroma); incident intensities of BDX and/or UVX were 1-2mW/mm^2^ (set to higher-end if ocular media was opaque), and/or ∼0.8mW/mm^2^ respectively. For far-red indicators, pre-mounted cube sets (Chroma 49015 and/or 89402) were used to image FR-GECO1c and/or other far-red indicators, and the incident intensity of RLX was ∼20-25mW/mm^2^. For violet illumination on top of constant blue illumination (Supplementary Fig. 1c): BDX incident intensity was 2mW/mm^2^ (10% X-Cite power) while UVX incident intensity was ∼0.8mW/mm^2^ (5% X-Cite power). UVX LED was turned on for 15s every minute by an external sequence generated by a Power3 DAQ and Spike2 software (Cambridge Electronic Design).

#### Imaging with a two-photon microscope

The microscopy setup was similar to (Barson et al., 2020). In brief: a movable objective two-photon microscope with a Janelia wide-path design and galvo-resonant scanner (Sutter Instruments) was used for *in situ* retina imaging. 920 nm infra-red laser was generated by a Ti:Sapphire laser (MaiTai eHP DeepSee, Spectra-Physics). An ultra-long-WD objective with a 20 mm WD and 0.40 NA (M Plan Apo NIR ×20, Mitutoyo) was used to image retina transpupillarily (retina was ∼2- 2.5mm below the cornea surface, depending on surgery and age conditions). Incident laser power was set to 50-100mW, depending on the opacity of the ocular media. Movies were acquired at 58.8Hz with bidirectional scanning and an averaging factor of 6 at resolution 256×256, rendering an effective frame rate of ∼10 Hz.

#### AAV-mediated expression of green or red indicators

Eye injections were conducted at P0-P1, and 500nL of AAVs (diluted to 1e13 GC/mL) was injected intravitreally into left eyes to express various calcium indicators. The injection protocol is similar to that described in (Gribizis et al., 2019a). 1. jGCaMP7 was expressed via AAV2.1-syn-jGCaMP7b-WPRE (Addgene, #104489-AAV1). 2. FR-GECO1c was expressed via AAV2.9-SYN-FR-GECO1c (Neurophotonics Centre, Lot #2556). HaloCaMP1b was expressed via AAV2.9-syn-HaloCaMP1b-EGFP (Addgene, #138328-AAV9), and Janelia Fluor® 635 (Tocris, Cat. No. 6419) was injected intravenously at a dose of 100 nmol 2 hours prior to imaging (Grimm et al., 2017) . 4. iGECI was expressed via AAV2.1-CAG-iGECI (generated by subcloning the Addgene plasmid #160423). 5. NIR-GECO2G was expressed via AAV2.1-CAG-NIR-GECO2G-WPRE (AAV generated by Yale Vision Core based on Addgene plasmid #159605). For positive control experiments, the same AAVs were injected to express corresponding indicators in cortex vias sinus injection (2uL on each side, diluted to 1e13 GC/mL) at P0-P1 (Hamodi et al., 2020).

### Data Analysis

#### Image pre-processing (Fig. 1-2 and related supplementary figures)

Raw TIFF movies were pre-processed with an object-oriented pipeline (https://github.com/CrairLab/Yixiang_OOP_pipeline). Parallel computing was realized in MATLAB on Yale’s high-performance clusters (Yale Center for Research Computing). The pipeline included the following steps: 1. Downsampling (average-pooling with a 2×2 filter). 2. Motion correction/subpixel rigid registration (Pnevmatikakis and Giovannucci, 2017) ROI mask application (manually defined in ImageJ prior to pre-processing) 4. Photobleaching correction using a single-term exponential fit was conducted on mean fluorescent traces of tiling 4×4 squares (e.g., for images of size 400 x 400, exponential fits would be conducted (400/4) x (400/4) = 10,000 times for all 4×4 squares tiling the field of view). 4. Gaussian smoothing (sigma/filter size = 1). 5. Denoising based on singular-vector decomposition (preserving top 256 dimensions) 6. Intensity normalization (ΔF/F, where F0 was defined as the 5th percentile value for each pixel). Key parameters were kept consistent across conditions.

#### Analysis of wave properties (Fig. 2 and Supplementary Fig.4)

Retinal wave properties were extracted by an integrated wave property analysis pipeline: WAIP (https://github.com/CrairLab/WAIP) that was modified based on code used in (Ge et al., 2021a). In brief: 1. pre-processed ΔF/F movies were z-scored and then binarized by threshold = 2 (consistent across conditions) 2. Connected components in the input binarized movies were analyzed with a MATLAB built-in function regionprops to extract a set of properties including duration, diameter, centroid, area etc. Connected components were filtered based on empirical thresholds to throw away components attributable to motion artifact/ noise (duration filter = 8 frames, diameter filter = 10 pixels). Only valid pixels were preserved for further analysis. 4. Vector fields Propagation directions of valid waves were extracted by a built-in MATLAB function estimateFlow based on the Lucas-Kanade method (Baker and Matthews, 2004). 5. Wave propagation directions were computed based on extracted vector fields. Key parameters were kept consistent across conditions. Wave property data was first combined across multiple movies of the same animal and then combined across animals of the same experimental group. As movies were acquired at various magnifications, all wave diameters were rescaled to the magnification level = 100x.

#### Opsin activation model (Supplementary Fig. 2)

Relative activations of mouse *rhodopsin in vivo* given different illumination conditions were estimated. First, spectral sensitivity was modeled assuming the maximum absorbance of mouse rhodopsin at λmax = 499 nm. Opsin template was adopted from equation 2 in (Lamb, 1995b):

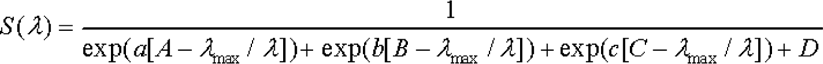

of which A = 0.880, B = 0.924, C = 1.104, D = 0.655, a = 70, b = 28.5, c = −14.1, and λ ranging from 350-1050nm.

To model the transmittance of mouse ocular media, equation 3 from (Lei and Yao, 2006a) was used to compute the spectral attenuation coefficient:

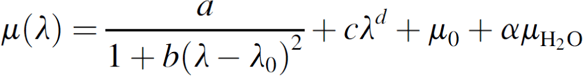

with values of model coefficients defined in Table 2 in the same paper.

Light percentage transmission of bandpass filters was acquired from the manufacturer Chroma (Chroma.com). X-Cite LED spectral intensities were estimated based on data provided by the manufacturer, assuming distributions were Gaussian with means and standard deviations estimated based on curve fitting results. Incident intensities were translated to photon flux (normalized to 460nm). Two-photon laser spectral intensities were estimated based on (Voigt et al., 2017). The overall activation of mouse rhodopsin under a specific condition was computed by integrating the above-mentioned factors below:

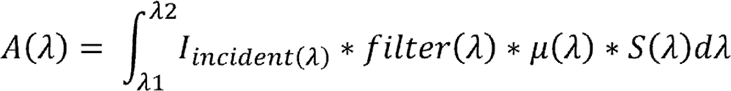

All parameters/ coefficients used in the model and the source code are available at https://github.com/CrairLab/Wave_model/blob/master/BestIndicator.m. Relative activations were computed with respect to the level of activation under the single-photon BDX illumination (5% X-CITE LED power) condition.

#### Seed-based correlation analysis

The number of reference seeds was pre-defined before parallel processing. Movies with a down-sampled field of view were densely covered by 500 evenly spaced seeds. Regular or partial Pearson-correlation matrices were generated with seeds inside ROI masks using built-in MATLAB functions “corr” and “partialcorr” respectively. For partial-correlation matrices, correlations between reference seeds and other pixels in the ROI were controlled for the averaged fluorescence trace over all pixels outside the ROI (to regress out the influence of non-specific whole-brain fluctuations). All seed-based correlation maps were manifested with jet (256) colormap and [-0.2, 1] color limit. Frames identified as containing motions (when amplitudes of rigid subpixel registration were larger than 0.5 pixel) were excluded using motion correction information generated in pre-processing procedures.

GraphPad Prism (RRID:SCR_002798) and Adobe Illustrator (RRID:SCR_010279) were used to generate/organize figures.

#### Motion energy

Motion energy was computed by 1. Summing pixel-wise changes in intensity between neighboring frames (captured by the infrared camera) 2. Normalizing frame-to-frame intensity illuminance by the total accumulated intensity change of the movie. 3. Denoising by a moving-average filter with a numerator coefficient inversely proportional to frame rates. The infra-camera was kept at the same angle and focus with a field of view including the front and rear limbs and the body of the animal.

#### Correlation between motion energy and calcium activity

Each session lasted for 2100 seconds (35 minutes), but only the last 30 minutes are used for analyses (the first 5 minutes were used to allow retina adaptation to the imaging scheme). Body movement movies and retina imaging movies (interleaved frames) were pre-processed and compared using the ReMo pipeline (https://github.com/CrairLab/ReMo). In brief: luminance differences of neighboring body-movement frames were computed, normalized, and filtered by an impulse response (FIR) filter (see function ComputeMotionEnergy). For the mesoscopic imaging data, interleaved frames were separated into blue and violet channels and processed separately. Each channel was downsampled, registered using NoRMCorre (Pnevmatikakis and Giovannucci, 2017), and photobleaching corrected. Processed violet-illuminated isosbestic signals were then regressed out from corresponding blue-illuminated signals to remove hemodynamic and motion artifacts (see function MotionActivityPreProcessing). The corrected mesoscopic ΔF/F data was then aligned with the motion energy trace. **To compute cross-correlations between corrected mesoscopic ΔF/F data and motion energy**: 1. Top-hat filtering was performed on mean ΔF/F (of the whole retina) with a hat = 3000 (frames). 2. Mean ΔF/F and motion energy trace were z-scored and smoothed with a filter of width 50 (frames). Cross correlations between the two processed traces were computed at lags up to ±1000 frames (±100s). Cyclic permutations were performed by cyclically shifting the motion energy trace by random intervals 500 times. **To plot time-lagged correlation maps:** 1. Top-hat filtering was performed on ΔF/F traces pixel-wise with a hat = 3000 (frames). 2. Top-hat filtered ΔF/F traces and motion energy trace was z-scored and smoothed with a filter of width 50 (frames). Pierson correlations were computed pixel-wise between individual ΔF/F traces and the motion energy trace. Correlations and p-values were reshaped to 2D maps.

## Author Contributions

Conceptualization, Y.W., and M.C.C.; Methodology, Y.W., D.B.; Software, Y.W.; Formal Analysis, Y.W.; Investigation, Y.W., A.S., J.J.; Resources, M.C.C.; Writing–Original Draft, Y.W.; Writing–Review & Editing, Y.W., J.J., M.C.C.; Visualization, Y.W.; Supervision, M.C.C; Project Administration, Y.W., M.C.C.

---

**Supplementary Fig. 1.**
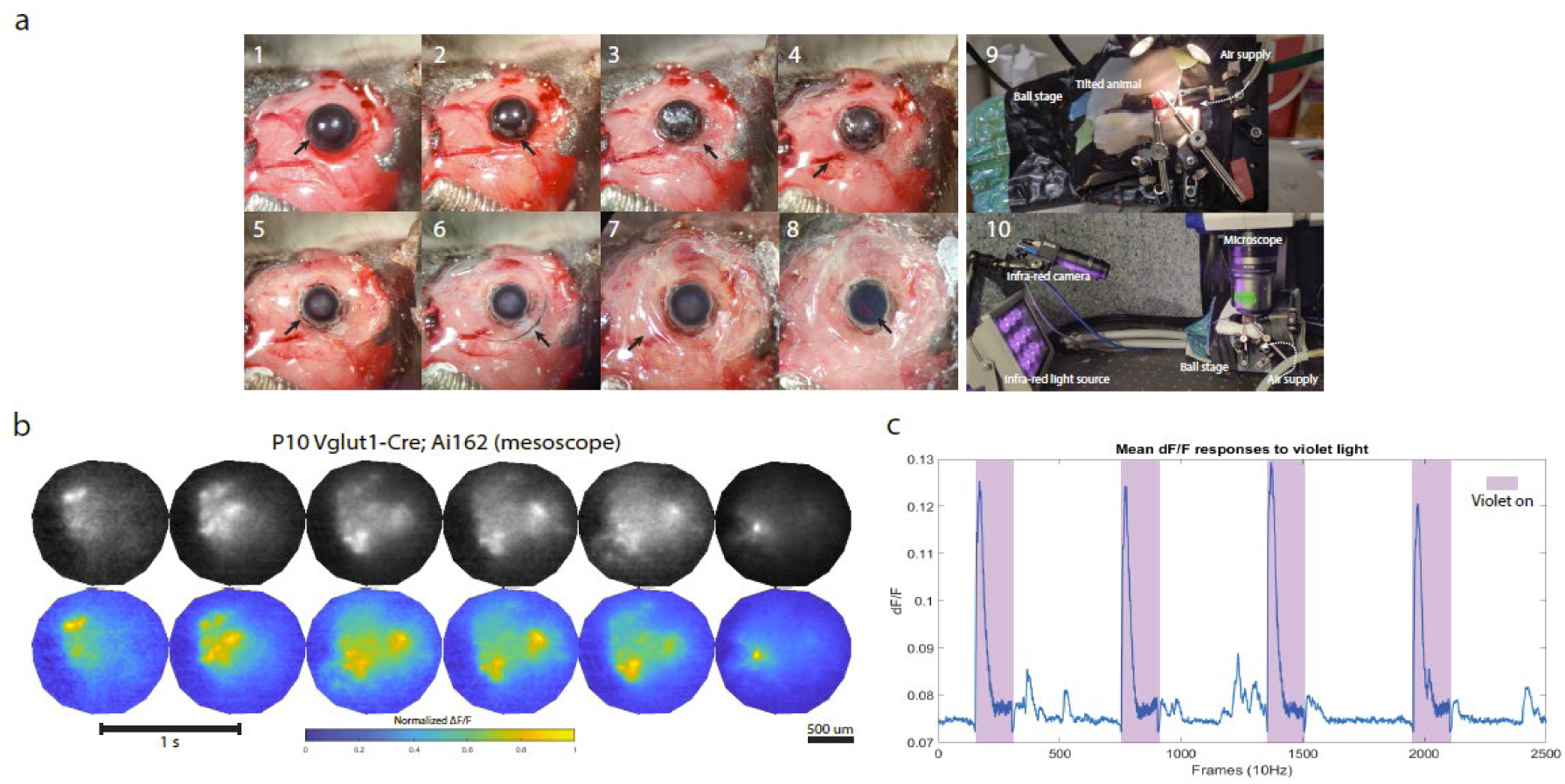
Eye surgery procedure (related to Fig. 1) a. Surgical procedure to stabilize eyeballs and the typical imaging configuration. Black arrows point to 1) an intact eyeball with eyelid and surrounding skin removed; 2) the eyeball after severing extra-ocular muscles, 3) dried Vetbond around the eyeball after an injection behind the eye, 4) Solidified cyanoacrylate covering the para-ocular area, 5) cleaned eye surface before hypromellose application, 6) 5mm coverslip floating on 2.5% hypromellose, 7) coverslip glued to surrounding skull with eyeball submerged in hypromellose underneath, 8) cleared lens and dark-purple reflection from the back of the eye after 90 min recovery from anesthesia, 9) Tilted ball stage during the surgical procedure, and 10) imaging configuration with an upright wide-field microscope and aninfrared camera to capture mouse body motions. b. Example montages of mesoscopic retina activity (retinal waves) from a P10 Vglut1-Cre; Ai162 animal. Colormaps: Montage interval = 1 second. ΔF/F values were normalized to [0, 1] for plotting. Top row: grayscale (raw fluorescence); Bottom row: parula (MATLAB). c. Example mean ΔF/F trace of mesoscopic retina activity from a P11 Snap25-GCaMP6s animal showed that retina could still respond to violet light (peaked at ∼385nm, incident intensity at the cornea surface: <1mW/mm^2^) on top of constant blue illumination (peaked at ∼460nm, incident intensity at the cornea surface: ∼2mW/mm^2^). Purple shades denote frames where violet light was on (every 60s for 15s). Note that: retinal waves were still present in between sessions (smaller peaks).

**Supplementary Fig. 2.**
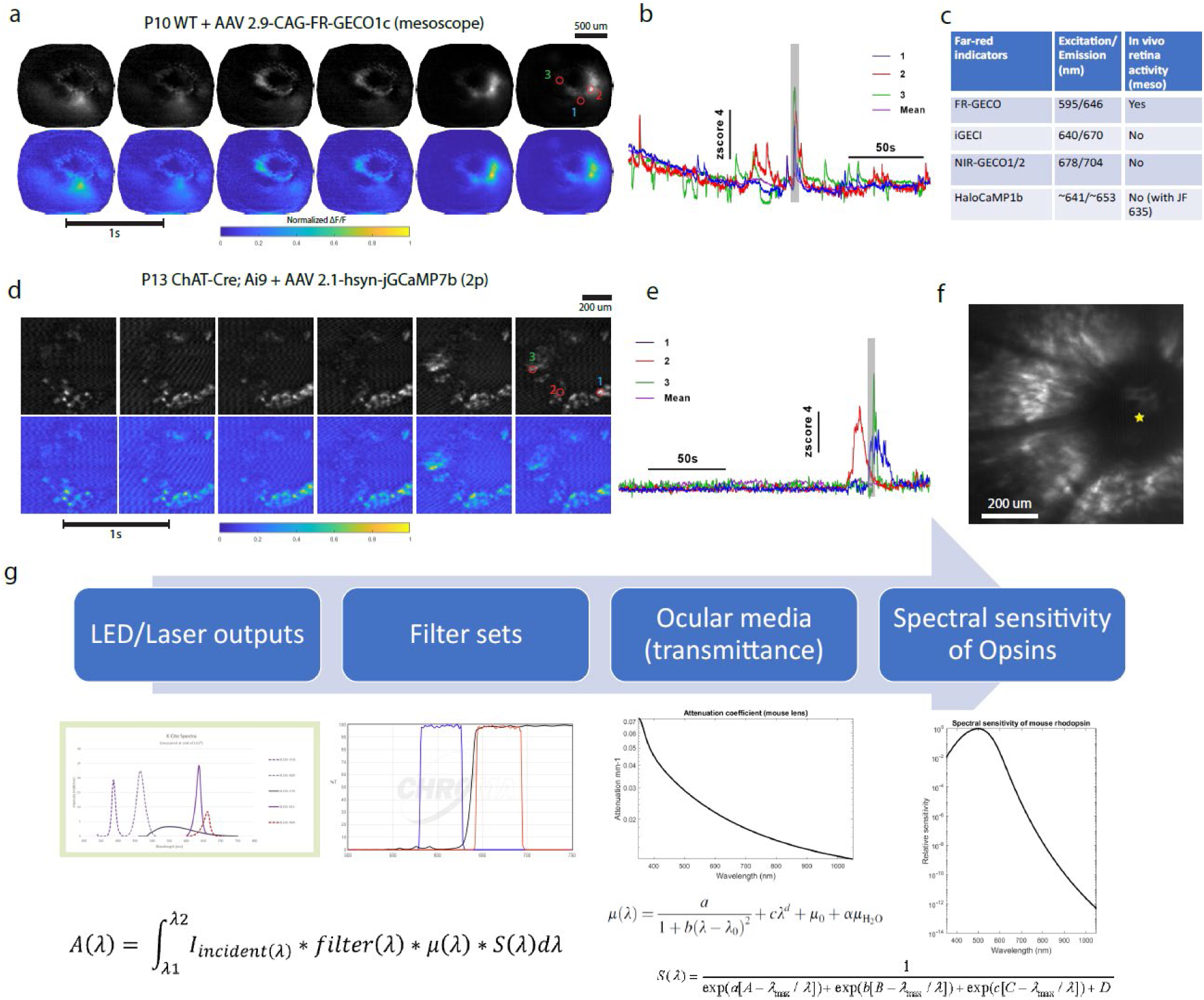
Imaging retina dynamics with far-red/infrared indicators (related to Fig. 1) a. Single-photon imaging with the FR-GECO1c indicator. Example montages of mesoscopic retina activity (retinal waves) from a P10 C57BL/6J animal (eye injection of AAV2.1-CAG-FR-GECO1c at P0). ΔF/F values were normalized to [0, 1] for plotting. Top row: grayscale (raw fluorescence); Bottom row: parula (MATLAB). Red dashed circles denote three pixels corresponding to the activity traces in panel b. b. Z-scored activity traces from three pixels/ ROIs denoted in panel a. Shaded area corresponds to period depicted in a. c. Table of far-red/infrared indicators tested. Indicators were expressed through eye injections of AAVs (see Method). Only FR-GECO1c showed detectable mesoscopic signals in retina imaging. All indicators except HaloCaMP1b showed detectable mesoscopic signals in cortex (see Method, data not shown). d. Two-photon imaging with jGCaMP7b. Example montages of spontaneous retina activity from a P13 ChAT-Cre; Ai9 animal (eye injection of AAV2.1-hsyn-jGCaMP7b at P0). ΔF/F values were normalized to [0, 1] for plotting. Top row: grayscale (raw fluorescence); Bottom row: parula (MATLAB). Red dashed circles denote three pixels corresponding to the activity traces in panel e. e. Z-scored activity traces from three pixels/ ROIs denoted in panel d. Shaded area corresponds to the period depicted in e. f. An example image of the field of view (FOV) generated by averaging over 4000 frames. Yellow star: optic disc. g. Workflow of modeling relative activation of mouse rhodopsin under different illumination conditions. Panels from left to right: 1) power spectra of different LEDs (translated to photon flux in the model), 2) transmittance curves of different filter sets, 3) attenuation curve of mouse ocular media (lens), and 4) relative spectral sensitivity of mouse rhodopsin. Formulas from left to right are for computing: 1, activations (A), 2) attenuation coefficients (u), and 3) relative spectral sensitivity (S).

**Supplementary Fig. 3.**
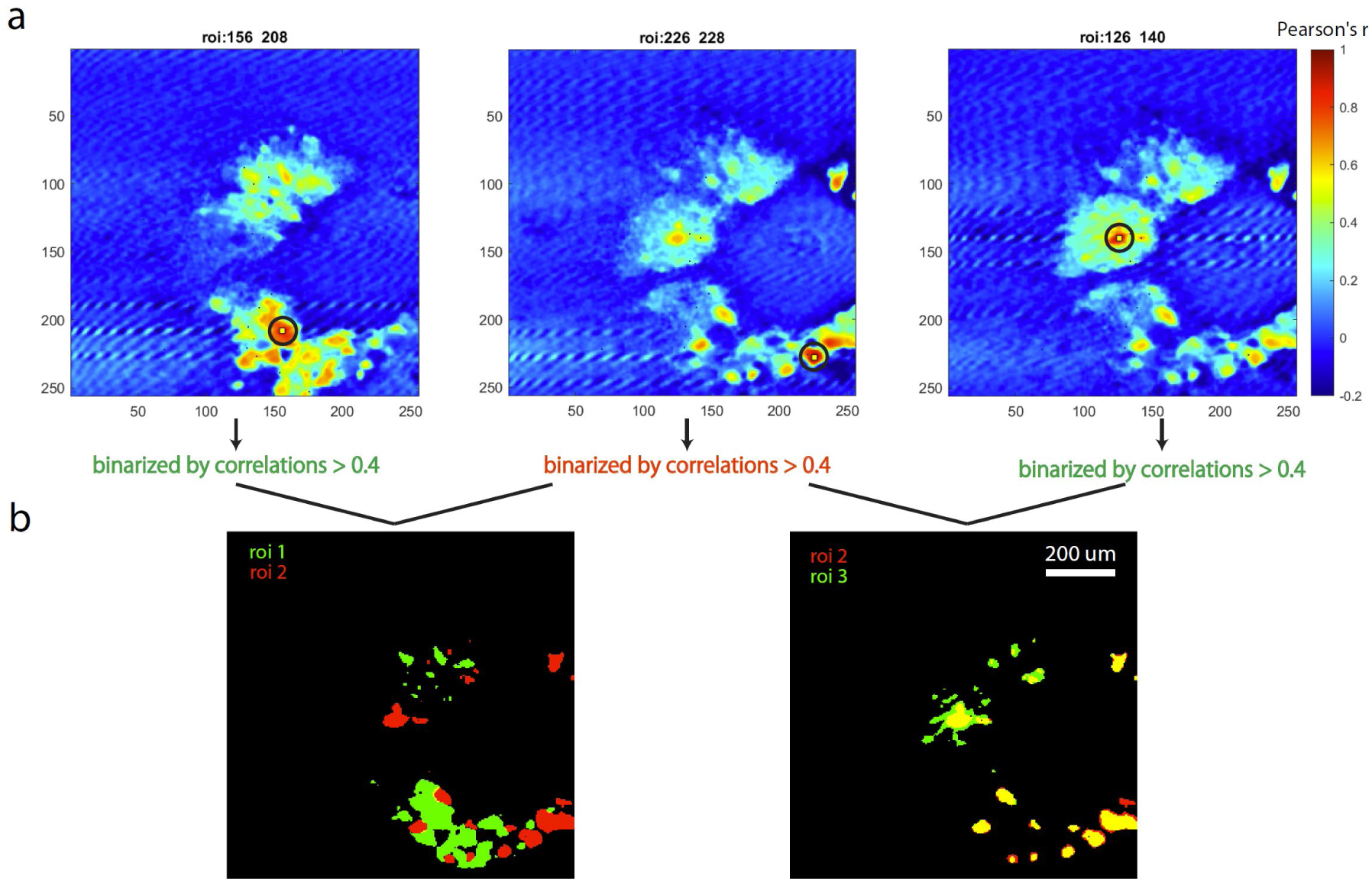
Seed-based correlation maps uncover distinct functional domains in the retina (related to Fig. 1) a. Seed-based correlation maps of three different ROIs (numbers indicate coordinates of reference seeds) based on two-photon imaging data. Yellow squares denote seed locations. Same data as in Supplementary Fig. 2d-f. b. Left: pseudo-colored merged image shows minimal overlap (<3%) between correlated regions (r > 0.4) of seeds 1 & 2 (green & red); right: substantial overlap (>80%) between correlated regions (r > 0.4) of seeds 2 & 3 (red & green). Overlapping area is shown in yellow.

**Supplementary Fig. 4.**
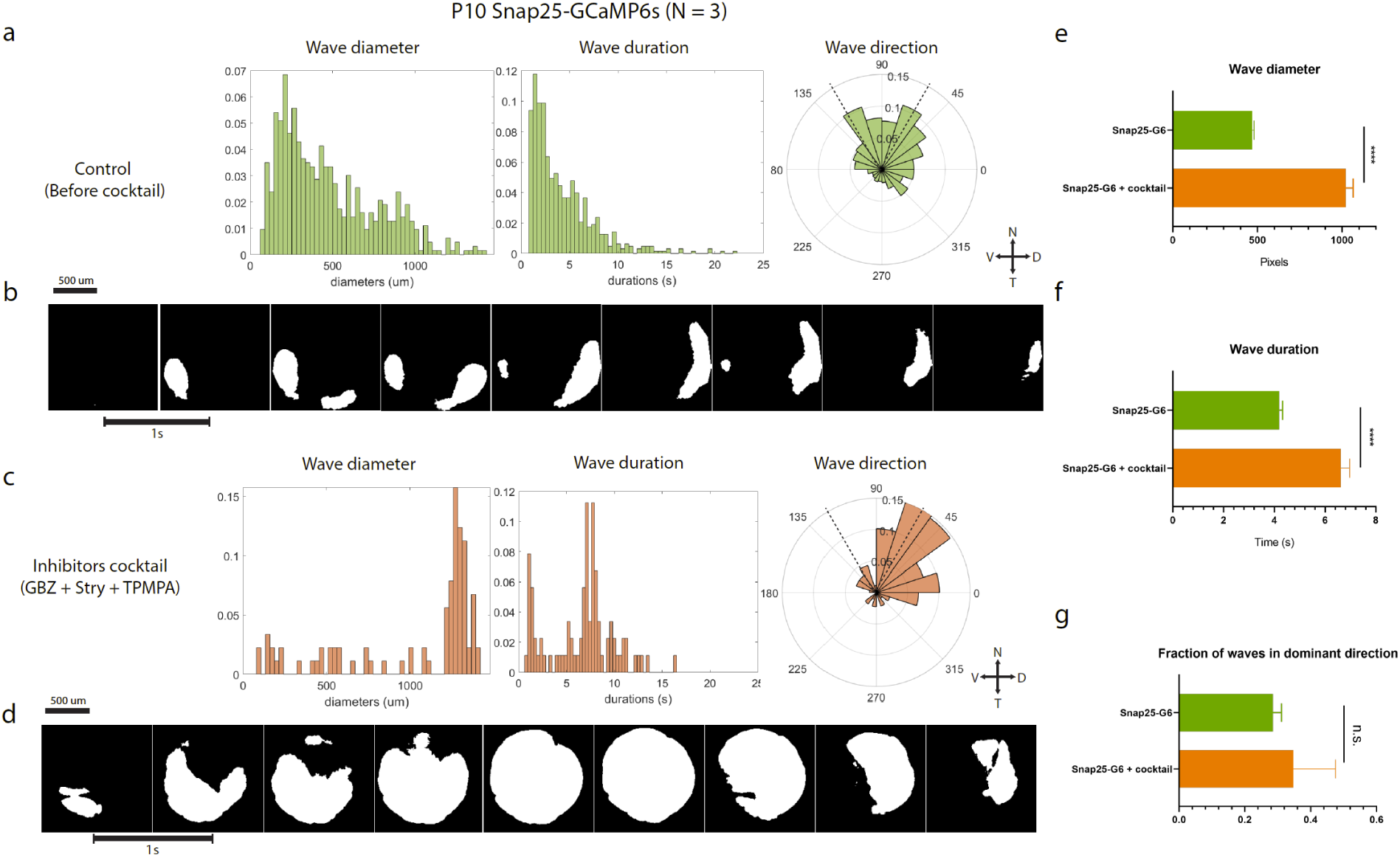
Antagonist cocktail altered spatiotemporal properties of stage III *in situ* retinal waves (related to Fig. 2) a. Spatiotemporal properties of stage III waves in P10-P11 SNAP25-GCaMP6s animals before application of antagonist cocktail (N = 3). Left: wave diameter distribution (normalized histogram, x-axis unit: um); Middle: wave duration distribution (normalized histogram, x-axis unit: seconds); Right: frequency distribution of waves directions (normalized polar histogram), 0 degree corresponds to the dorsal direction. Crossed double-headed arrows indicate cardinal directions (N: nasal; V: ventral; D: dorsal; T: temporal). Dashed black line, fraction in the dominant direction (temporal-nasal) if wave propagation directions are random. b. Montages of example retinal waves (binarized, z-score threshold = 2) in control. Active pixels are in white. c. Spatiotemporal properties of stage III waves in P10-P11 SNAP25-GCaMP6s animals after application of cocktail solution (N = 3). 250-300nL of cocktail solution (200uM Gabazine + 100uM Strychnine + 2mM TPMPA) was injected intravitreally (see Method for details). Panel layout similar to a. d. Montages of example retinal waves (binarized, z-score threshold = 2) after cocktail solution. Active pixels are in white. e. Diameters of retinal wavers were significantly larger after cocktail application. f. Duration of retinal waves was significantly longer after cocktail application. g. Proportion of wave propagation directions within 60° surrounding the dominant direction (temporal-nasal) did not show significant change before/after the cocktail. n.s. p > 0.05, ∗ p < 0.05, ∗∗ p < 0.01, ∗∗∗ p < 0.001, **** p < 0.0001, two-tailed unpaired t-test with Welch’s correction.

**Supplementary Fig. 5.**
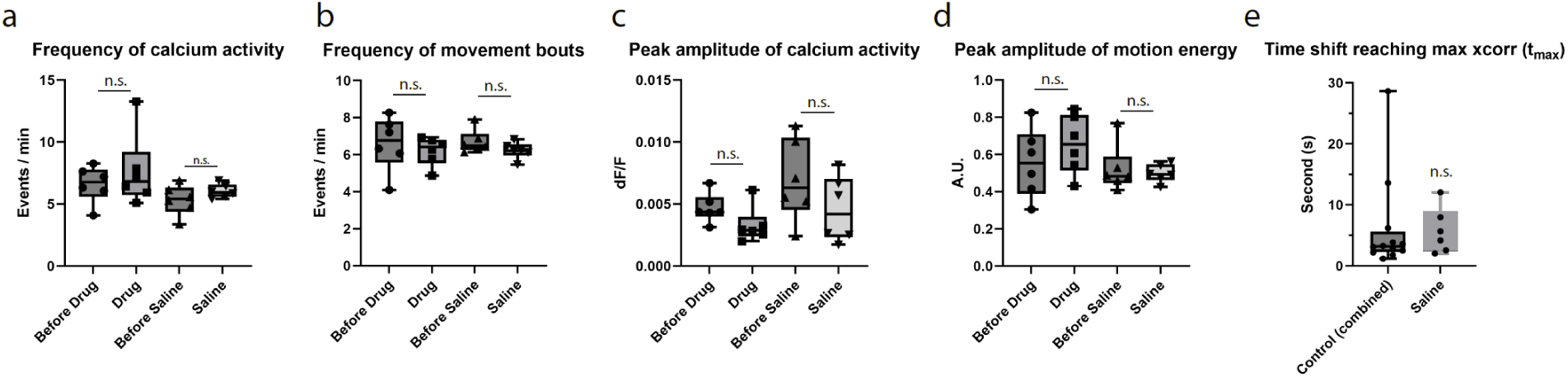
Properties of calcium activity and movement under different conditions (related to Fig. 3) a. Frequency of mean calcium activity before/after mepyramine injection and before/ after saline injection (quantified by identifying ΔF/F peaks in average calcium traces of all pixels in the field of view, N = 6 for each group, same animals as in Fig. 3.) b. Frequency of movement bouts before/after mepyramine injection and before/after saline injection (quantified by identifying motion energy peaks). c. Peak amplitude of mean calcium activity before/after mepyramine injection and before/after saline injection (quantified by the prominence of ΔF/F peaks) d. Peak amplitude of movement bouts before/after mepyramine injection and before/after saline injection (quantified by the prominence of motion energy peaks.) e. Time shifts (positive t_max_ indicates the amount of time that calcium activity trailing movement) where max cross-correlations were found. Control (combined) group: N = 12 (average tmax = 6.1s, median tmax = 3.2s), combing 6 animals before mepyramine injection and 6 animals before saline injection. Saline group: N =6 (average tmax = 5.7s, median tmax = 4.9s) n.s. p > 0.05, ∗ p < 0.05, ∗∗ p < 0.01, ∗∗∗ p < 0.001, **** p < 0.0001, two-tailed unpaired t-test with Welch’s correction. For all Box plots in this study: hinges: 25 percentile (top), 75 percentile (bottom). Box whiskers (bars): Max value (top), Min value (bottom). The line in the middle of the box is plotted at the median.

**Supplementary Fig. 6.**
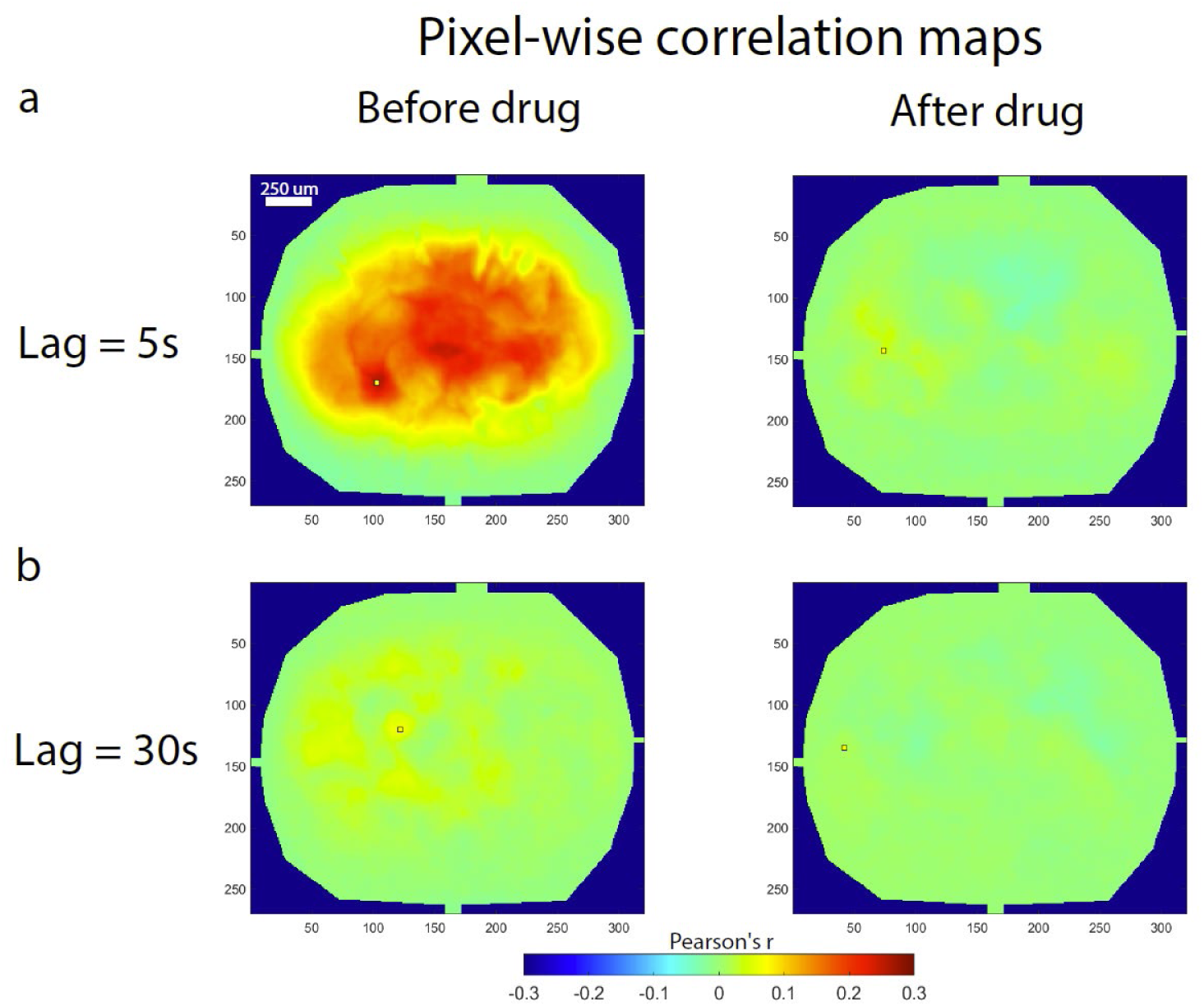
Cross-correlation and pixel-wise correlation maps in high/low calcium activity groups (related to Fig. 3) a. All control data (N = 12, merged from 6 animals before mepyramine injection and 6 animals before saline injection) were regrouped into a high-activity cohort (mean peak ΔF/F ranging from 0.0052-0.0113, N = 6) and a low-activity cohort (mean peak ΔF/F ranging from 0.0024-0.0052, N = 6). Cross-correlation and pixel-wise correlation map analyses (averaging across 6 animals in each cohort) were conducted to show changes in calcium activity level (ΔF/F amplitude) and did not affect correlational results.

